# GIP: An open-source computational pipeline for mapping genomic instability from protists to cancer cells

**DOI:** 10.1101/2021.06.15.448580

**Authors:** Gerald F. Späth, Giovanni Bussotti

**Affiliations:** Institut Pasteur, INSERM U1201, Unité de Parasitologie moléculaire et Signalisation, Paris, France; Hub de Bioinformatique et Biostatistique - Département Biologie Computationnelle, Institut Pasteur, 75015 Paris, France

**Keywords:** Genome instability, aneuploidy, copy number variations, single nucleotide variation, structural variation, genetic adaptation

## Abstract

Genome instability has been recognized as a key driver for microbial and cancer adaptation and thus plays a central role in many human pathologies. Even though genome instability encompasses different types of genomic alterations, most available genome analysis software are limited to just one kind mutation or analytical step. To overcome this limitation and better understand the role of genetic changes in enhancing pathogenicity we established GIP, a novel, powerful bioinformatic pipeline for comparative genome analysis. Here we show its application to whole genome sequencing datasets of *Leishmania*, *Plasmodium, Candida*, and cancer. Applying GIP on available data sets validated our pipeline and demonstrated the power of our analysis tool to drive biological discovery. Applied to *Plasmodium vivax* genomes, our pipeline allowed us to uncover the convergent amplification of erythrocyte binding proteins and to identify a nullisomic strain. Re-analyzing genomes of drug adapted *Candida albicans* strains revealed correlated copy number variations of functionally related genes, strongly supporting a mechanism of epistatic adaptation through interacting gene-dosage changes. Our results illustrate how GIP can be used for the identification of aneuploidy, gene copy number variations, changes in nucleic acid sequences, and chromosomal rearrangements. Altogether, GIP can shed light on the genetic bases of cell adaptation and drive disease biomarker discovery.

**One Sentence Summary:** GIP - a novel pipeline for detecting, comparing and visualizing genome instability.

## Main Text

In recent years, the field of genomics has rapidly expanded with a fast increase in the number of newly sequenced genomes (1). This surge is a direct consequence of the development of new and ever more efficient, high-throughput capable sequencing technologies (2). On the one hand, the improvement in long reads technology allowed the generation of high-quality genome assemblies (3,4). On the other hand, the decreasing costs for short-reads sequencing and the parallel increase in sequencing throughput propelled the exponential increase of available whole genome sequencing (WGS) data (5). Thanks to these advances one can reasonably expect WGS to rapidly become a key component of personalized medicine and clinical applications. In this context several consortia-based projects have been established with the goal to produce WGS for the study of different biological systems (5), and a number of publicly available databases have been compiled or updated (6). Parallel to data availability, many bioinformatics tools have been developed to perform specific genome analysis tasks (7,8). For instance, tools such as Freebayes (9), CNVnator (10) and DELLY (11) have been respectively used for the detection or characterization of DNA single nucleotide variants (SNVs), copy number variations (CNVs), and structural variations (SVs), but their scope is limited to the analysis of one genomic feature at the time. A number of integrative WGS pipelines and workflows have been established combining the execution of multiple bioinformatics algorithms serving different analysis steps (12). Even though continuous progress has been made, there is no standardized or unified approach for genomic investigation. For the development of improved WGS data analysis pipelines, several important requirements need be considered, including portability, reproducibility, scalability and compatibility with high-performance computing (HPC) clusters and remote cloud computing. Here we introduce a novel genome instability pipeline (GIP) that fulfills all these requirements. GIP facilitates the genome-wide detection, quantification, comparison and visualization of chromosome aneuploidies, gene CNVs, SNVs and SVs. GIP is implemented in Nextflow (13), a workflow language that allows to execute GIP seamlessly in local workstation, on an HPC or remotely in the cloud. All required environment and software dependencies of GIP are fulfilled and provided with a Singularity container, thus making GIP reproducible, easy-to-install and easy-to-use. GIP allows the use of giptools, a tool-suit of R-based modules for genome data exploration, enabling the comparison of sample sub-sets. GIP and giptools generate a summary report with publication-quality figures and spreadsheet tables. GIP and giptools constitute a single framework for WGS analysis suitable both for large scale batch analysis of individual genomes and comparison of samples from different experimental conditions or origins. Lastly, a key strength of GIP and giptools is the general applicability to different eukaryotic species. We already successfully applied GIP on the analysis of *Leishmania* genomes (14–16). In this study, we validate the use of GIP and giptools using WGS data from published datasets of the three major human pathogens *Leishmania infantum*, *Plasmodium vivax* and *Candida albicans* and as well as three human cancer cell lines. Furthermore, we demonstrate how the extensive and powerful analytical approach operated by GIP and giptools can be used to find new biological signal that escaped previous analyses.

## Results

### The GIP workflow

GIP is a tool for scientific investigation compatible with Linux systems, requiring minimal configuration and distributed as a self-contained package. GIP consists of three files: the Nextflow pipeline code, the configuration and the Singularity container files. The minimum required input is a paired-end WGS data set and a reference genome assembly in the standard fastq and FASTA format, respectively. GIP analyses include (i) extracting genomic features such as assembly gaps or repetitive elements, (ii) mapping the reads, (iii) evaluating chromosome, gene and genomic bin copy numbers, (iv) identifying and visualizing copy number variation with respect to the reference genome, (v) identifying and quantifying gene clusters, (vi) detecting and annotating SNVs, (vii) measuring non-synonymous (N) and synonymous (S) mutations for all genes, (viii) detecting SVs including tandem duplications, deletions, inversions and break-ends translocations using split-read and read-pair orientation information, and (ix) producing a report file providing summary statistics, tables and visualizations (**Fig. 1**). GIP allows to customize filtering and visualization options via the configuration file (see methods). The output of GIP can be used as input for giptools, a tool-suite to compare sample sub-sets and highlight chromosome copy number, gene copy number and SNV differences.

**Fig. 1.**
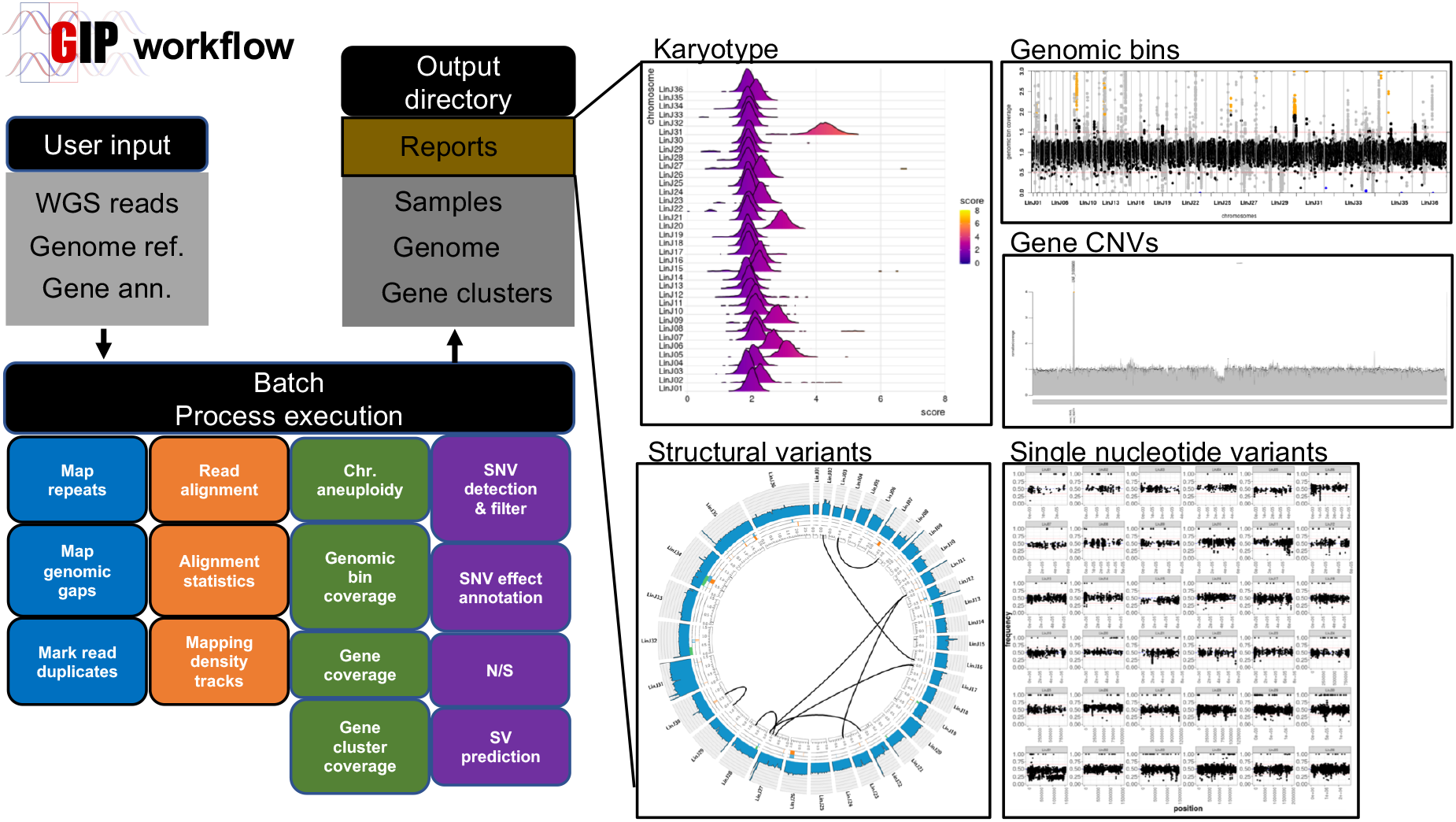
GIP workflow. The schema on the left recapitulates the GIP inputs, processes and outputs (see methods). Blue, orange, green and purple boxes indicate genome reference, read mapping, quantification and variants computation steps, respectively. The panels on the right demonstrate example plots included in the GIP report computed for individual samples. The “Karyotype” plot shows the coverage distributions for each chromosome (y-axis). The “Genomic bins” plot shows the genomic position (x-axis) and the normalized genomic bin sequencing coverage (y-axis). The “Gene CNVs” plot shows the normalized gene sequencing coverage. The “Structural variants” panel shows a Circos plot representing break-end translocations (black links in the inmost part of the plot), and other possible structural variations in the outer tracks, including insertions, duplications, deletions and inversions. The outmost track shows the normalized sequencing coverage. The “Single nucleotide variants” plot shows on the x and y axes respectively the genomic position and variant allele frequency of detected SNVs.

### Applying giptools on a *Leishmania infantum* case study

GIP permits the batch analysis of a set of individual samples, where each sample is considered separately and compared only with respect to the provided reference genome assembly. As a consequence, all variants and copy number alterations detected in a sample merely reflect the differences between the sequenced and the reference genomes. While this application may be sufficient in some circumstances, research projects often involve downstream comparison between samples. Examples include the comparison of gene or chromosome copy variation number between for example drug resistant and drug susceptible samples, or the juxtaposition of SNVs detected in isolates from different geographic areas. For this purpose, we developed giptools, a suit of thirteen modules that allows to compare samples processed by GIP (**Table 1**). All modules in giptools are fully embedded in the Singularity container and they are provided with their own documentation.

**Table 1:**
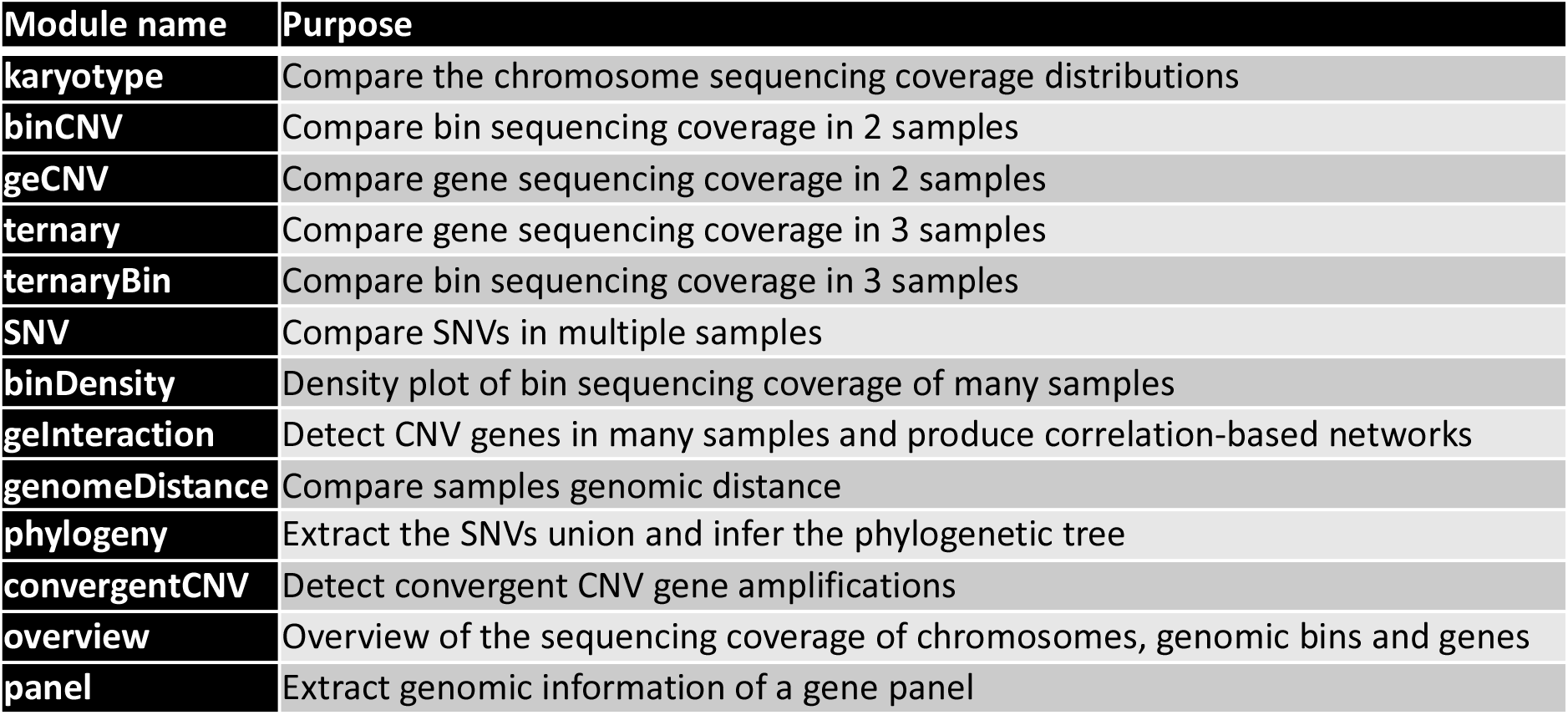
giptools modules.

To illustrate the type of exploratory data analyses and the biological questions that can be addressed, we tested giptools on a previously analyzed dataset of seven clinical *Leishmania infantum* isolates from Tunisia (14). *Leishmania* is the etiological agent of leishmaniasis, a life-threatening human and veterinary disease affecting 12 million people worldwide (17). The parasites were derived from patients affected with visceral leishmaniasis, expanded in cell culture and their genomes were sequenced. The dataset includes four Glucantime drug susceptible isolates and three isolates from relapsed patients, and thus comparison may inform on genetic factors resulting in treatment failure. giptools allowed the detection and visualization of pervasive intra-chromosomal CNVs across the thirty-six *Leishmania* chromosomes (**Fig. 2A**). Additionally, giptools enables targeted comparison of normalized genomic bin sequencing coverage of sample pairs. We used giptools’ ‘binCNV’ module to compute the ratio between corresponding genomic bins of the strains LIPA83 over ZK43, which correspond to a first-episode and a relapse leishmaniasis isolate, respectively. giptools represents different chromosomes as separate panels (**Fig. 2B**), as part of single genome-wide overview (**Fig. S1A**) or as distinct plots (**Fig. S1B**). This analysis allowed the identification of 2,905 and 2,208 bins that were respectively amplified or depleted in LIPA83 with respect to ZK43. The results are returned by giptools as a Microsoft Excel table (.xlsx format) providing ratio scores at each genomic position (**Table S1**). Likewise, giptools permits three-way comparisons of normalized genomic bin sequencing coverage with ternary plots (**Fig. 2C**). We used this representation to display the genomic bin relative abundance in samples ZK43, LIPA83 and ZK28. The analysis shows important strain-specific differences in bin copy number that are visualized by shifts of the signals out of the center. Similar to genomic bin analysis, giptools makes it possible to compare the sequencing depth of annotated genes and thus the copy number in two or three samples (**Fig. 2D** and **Fig. S2**). However, the determination of gene copy number might be impeded by (i) short read length or fragment insert size, (ii) the complexity of the target genome, or (iii) the presence of repetitive elements. The read map quality (MAPQ) score is a measure that reflects how much each gene is supported by unambiguously mapped reads (high MAPQ) in contrast to multimapping reads (low MAPQ). Together with the coverage, GIP also computes the mean read MAPQ score for each gene and allows a different strategy to determine the copy number of low MAPQ genes (see methods). The evaluation of LIPA83 and ZK43 gene coverage ratio scores revealed 13 gene CNVs with a stringent MAPQ cutoff of 50 (**Table S2**). The maximum normalized coverage value of 6.5 was observed for a putative amastin surface glycoprotein (LINF_310009800). Other examples of gene CNVs in this set include the putative surface antigen protein 2 (LINF_120013500) and the heat shock protein HSP33 (LINF_300021600) (**Table S2**). Genes falling below a user defined MAPQ score and sharing high level of sequence similarity are assigned to the same gene cluster, and their measured coverage scores are averaged across all members of the group. Low MAPQ scores can also be associated with single genes, e.g. in the case of internal repetitive elements that cause multiple ambiguous alignments inside the gene itself, or if mapping occurs in possibly misannotated intergenic regions. The LIPA83/ZK43 comparison showed 27 CNV gene clusters, including cluster clstr303 (3 genes annotated as ‘amastin-like’) and cluster clstr16 (2 tb-292 membrane-associated protein-like proteins) (**Table S2**). These results demonstrate the power of GIP and giptools to detect and compare intra-chromosomal CNVs in *Leishmania* at genomic bin level. Conveniently, analogous two- or three-ways comparisons can be applied to reveal copy number variations at individual gene or gene cluster levels.

**Fig. 2:**
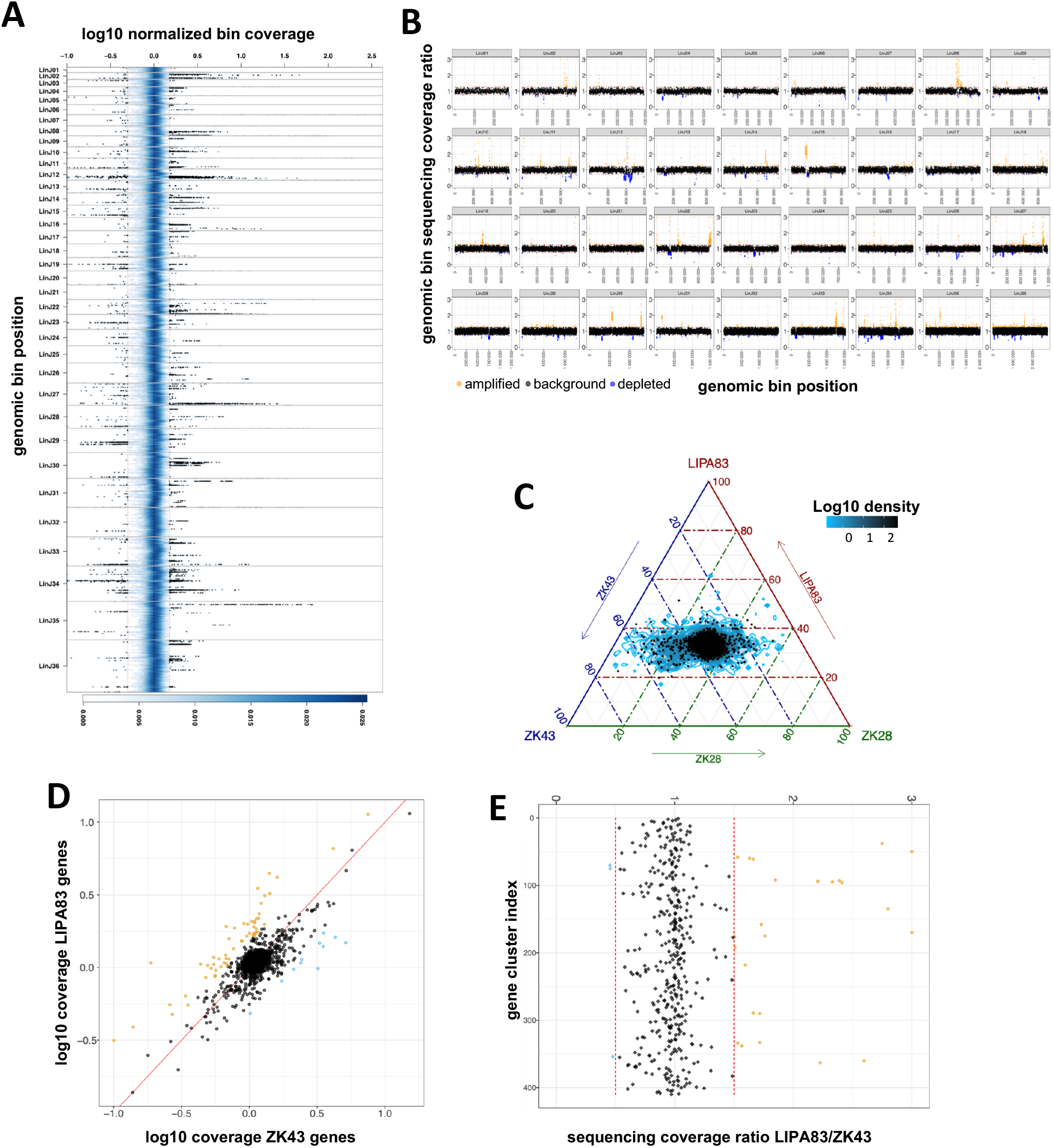
Comparing *Leishmania infantum* genomes with giptools. (**A**) Density plot representing the genomic coverage of the seven *Leishmania infantum* isolates. The x-axis shows the log 10 normalized coverage of genomic bins. The y-axis reflects the genomic position. The thirty-six different chromosomes are materialized as separate panels. The blue shading indicates the (2D) kernel density estimates of genomic bins. The two red vertical lines mark the 1.5 and 0.5 coverage values. A selection of 50 000 bins with coverage > 1.5 or < 0.5 is shown as black dots. (**B**) Scatterplot of the genomic bin normalized sequencing coverage ratio of samples LIPA83 over ZK43. The x any y axes show the ratio score and the genomic position respectively. Ratio scores > 1.25 are labelled in orange and indicate genomic bin amplification. Ratio scores < 0.75 are labelled in blue and indicate genomic bin depletion. (**C**) Ternary comparison showing the relative abundance of the genomic in samples LIPA83, ZK43 and ZK28. The axes report the fraction of the bins normalized sequencing coverage in the three strains. The blue contour indicates the log10 bin density. A subset of 5,000 bins is shown as black dots. Each given point in the plot is adding up to 100. The density area at the center of the plot indicates bins with equal copy number and thus a ~33 distribution across the three axes. (**D**) Scatterplot showing the log10 normalized sequencing coverage of annotated genes in ZK43 (x-axis) and LIPA83 (y-axis). The red line indicates the bisector. Dots represent individual genes. (**E**) Sequencing coverage ratio of gene clusters in samples LIPA83 and ZK43. Dots represent gene clusters. For plots **D** and **E**, the ratio scores > 1.5 or < 0.5 are labelled in orange and blue respectively.

### Comparative genomics of a *Plasmodium vivax* WGS dataset

We next applied GIP and giptools on other biological systems to demonstrate its broad applicability outside the *Leishmania* field, including the human apicomplexan parasite *Plasmodium vivax*. *Plasmodium vivax* is a protist parasite and a human pathogen causing malaria. *Plasmodium vivax* gives rise every year to 130 million clinical cases (18), and it is estimated that 2.5 billion people are at risk of infection worldwide (19–21). We applied GIP and giptools to investigate genomic variations across a sizeable dataset of 222 *Plasmodium vivax* genomes isolated from clinical samples of 14 countries worldwide (22,23) (**Table S3**). The phylogenetic tree reconstruction and PCA analyses (**Fig. 3A** and **B**) showed a high correlation between genotypes and the geographic origin of the samples. However, we detected substantial genomic variability between isolates collected at smaller geographical scale, with 14,555 SNVs (~42% of the total) uniquely characterizing representative samples from five Ethiopian study sites (23) (**Fig. 3C** and **D**). This result may reflect diverging evolutionary trajectories radiating from few founder strains. At gene level we profiled the copy number variations of two gene panels. The first panel accounts for 43 previously described genes encoding for potential erythrocyte binding proteins suggested to operate at the interface of the parasite-host invasion process (23). The second panel includes two drug resistance markers comprising the chloroquine resistance transporter PVP01_0109300 and the multidrug resistance protein 1 PVP01_1010900, and four proteins implicated in red blood cell invasion, such as the merozoite surface protein gene MSP7 PVP01_1219700, the reticulocyte binding protein gene 2c PVP01_0534300, the serine-repeat antigen 3 PVP01_0417000, and the reticulocyte binding protein 2b PVP01_0800700) (24–33). Read depth analysis indicated that four genes in the panel (PVP01_0623800, PVP01_1031400, PVP01_1031200, PVP01_1031300) show a high degree of variability, with amplifications observed in samples from distinct geographic (**Fig. 3E**). This convergence is sign of strong natural selection, which further sustains the functional importance of these genes in the infection process. Furthermore, 6 genes positioned on chromosome 14 are absent in the Thai strain PD0689_C as a result of the loss of this chromosome (nullisomy) (**Fig. 3E** and **F**). Finally, the comparison of synonymous and non-synonymous SNVs in the panel of genes revealed important differences between sample groups. Our analysis indicates an overall higher number of non-synonymous mutations in Ethiopian compared to Cambodian isolates, therefore suggesting a stronger evolutionary pressure acting on the African strains (**Fig. 3G**). Taken together these analyses well illustrate how GIP and giptools can be readily applied for bulk analysis of *Plasmodium vivax* genomes to assess genome diversity, extract evolutionary information and identify potential disease biomarkers.

**Fig. 3:**
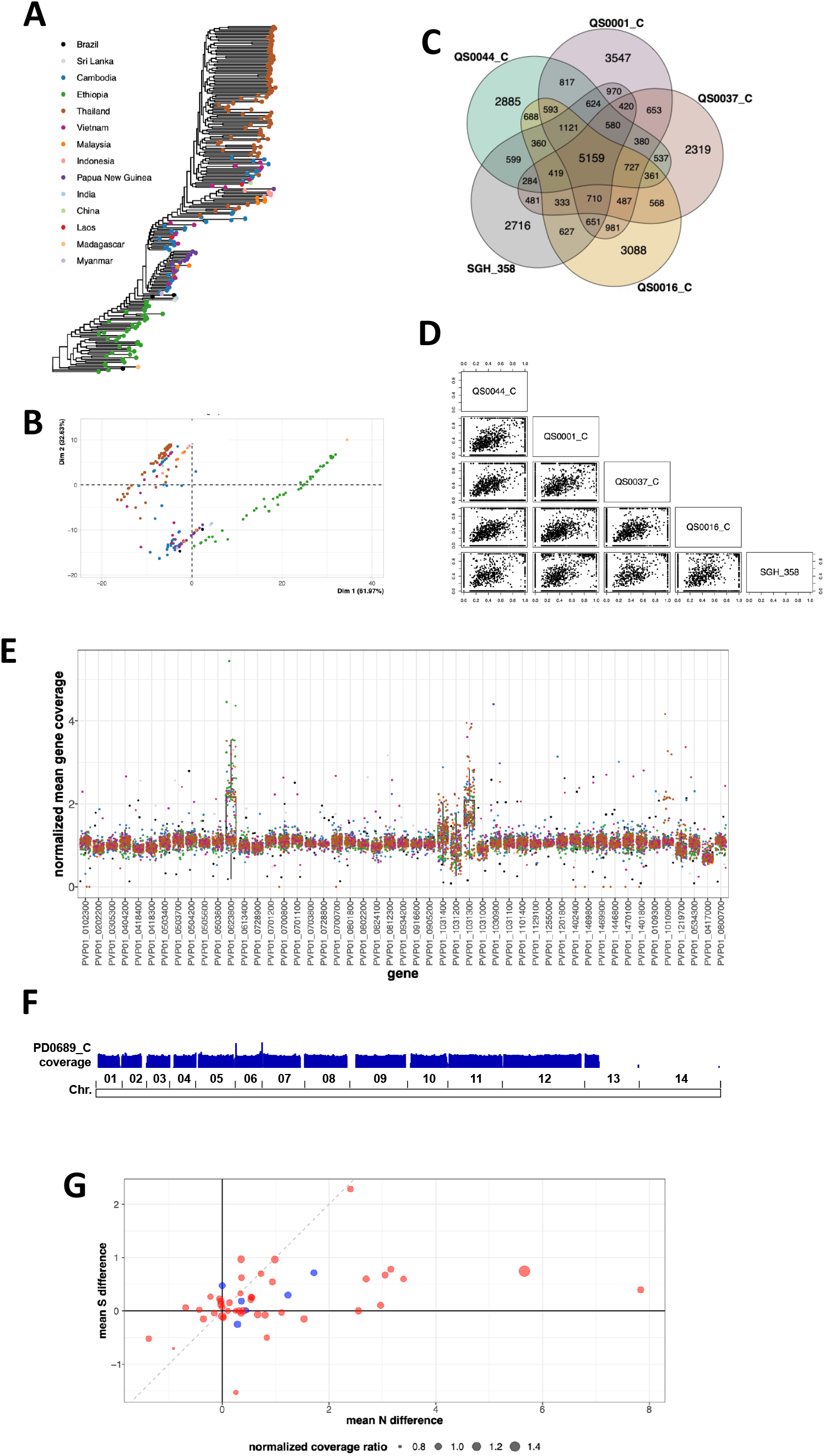
*Plasmodium vivax* genomic diversity. (**A**) Predicted maximum likelihood phylogenetic tree reconstruction. (**B**) PCA analysis of the phylogenetic distances estimated from the tree in (**A**). Each dot indicates a sample. The colour code reflects the geographic origin of the samples and matches with the colours of the legend in (**A**). (**C**) Venn diagram comparing the SNVs of five representative Ethiopian strains. (**D**) Pairwise scatterplot comparing the variant allele frequency of all detected SNVs in the five Ethiopian strains. (**E**) Gene panel analysis. The x-axis reports a set of genes of interest. The y-axis indicates the normalized mean gene coverage. The boxplots demonstrate the coverage values distributions for each gene across all samples. Each dot represents the coverage of the indicated gene in a given sample. Dot colours reflect the sample geographic origin as in (**A**). (**F**) Reads per kilo base per million mapped reads (RPKM) normalized sequencing coverage density track of sample PD0689_C. The boundaries of the 14 chromosomes are shown on the bottom. (**G**) Comparison of non-synonymous (N) and synonymous (S) mutations between Ethiopia and Cambodia sample groups. Dots represent genes. The x-axis represents the difference between the mean non-synonymous mutation count in the two sample groups. The y-axis represents the difference between the mean synonymous mutation count in the two sample groups. The dot size demonstrates the ratio of the mean normalized sequencing coverage between the two sample groups for each gene. Red and blue dot colors indicate genes belonging to the 43 genes panel (23) and the custom 6 genes panel respectively.

### Gene CNV analysis of *Candida albicans* evolutionary adapted strains

We next applied GIP and giptools to the human fungal pathogen *Candida albicans*, an opportunistic yeast exhibiting major genome plasticity (34–43) and causing hundreds of thousands of severe infections each year (44). Candidemia, a bloodstream infection with *Candida*, are often associated with high rates of morbidity and mortality (15–50%) notwithstanding existing antifungal treatments (45,46). We applied GIP and giptools to a *Candida albicans* WGS dataset described in a recent study that covers five different progenitor strains (P75063, P75016, P78042, SC5314, AMS3050) and investigates CNVs driving tolerance and resistance to anti-fungal azole drugs (47). We analyzed nineteen samples, including (i) four clinical isolates (P75063, P75016, P78042, SC5314), (ii) seven strains selected *in vitro* against the anti-fungal drug fluconazole (FLC) (AMS4104, AMS4105, AMS4106, AMS4107, AMS4397, AMS4444, AMS4702), (iii) four isogenic colonies adapted to the drug miconazole (AMS3051, AMS3052, AMS3053 and AMS3054) together with their progenitor (AMS3050), and (iv) three colonies derived from a miconazole-adapted population and isolated on a rich medium (AMS3092, AMS3093 and AMS3094) (47–49). GIP and giptools were able to reproduce previous observations of the amplification of the genes for the drug efflux pumps TAC1 (orf19.3188) and ERG11 (orf19.922), for the stress response proteins HSP70 (orf19.4980), CGR1 (orf19.2722), ERO1 (orf19.4871), TPK1 (orf19.4892), ASR1 (orf19.2344), PBS2 (orf19.7388) and CRZ1 (orf19.7359), and for proteins involved in membrane and cell wall integrity, including CDR3 (orf19.1313), NCP1 (orf19.2672), ECM21 (orf19.4887), MNN23 (orf19.4874), RHB1 (orf19.5994) and KRE6 (orf19.7363) (**Table S4**). Furthermore, the powerful comparative approach of our pipeline permitted the discovery of 1,505 genes showing correlating or anti-correlating copy number variations (**Fig. 4A** and **B, Table S4**), which could be assigned to nine distinct correlation clusters (CC) (**Fig. S3A, Table S5**) that escaped previous analyses. We verified the sequencing coverage of genomic regions encompassing gene CNVs, including three regions amplified in fluconazole resistant strains (**Fig. 4B, Fig. S3B**) (47) and a region whose amplification correlates with the level of miconazole resistance (47) (**Fig. S3C**), as well as the loss of heterozygosity associated to the depletion of chromosome 3 left arm in sample AMS3051 (**Fig. S3D**) (47). Eventually, by representing genes and absolute correlation respectively as nodes and edges of a network, we identified 9 highly interconnected network clusters (NC) (**Fig. 4C, Table S6**). NC7, NC8 and NC9 embody genes from individual chromosomes, respectively chromosomes 1, 3 and 4. The most parsimonious explanation for the high levels of correlation observed in these NCs (**Fig. 4C**) is the occurrence of sub-chromosomal amplifications affecting several adjacent genes. A different scenario is pictured for each of the remaining NCs (NC1-6) where the genes are located on different chromosomes thus suggesting genetic interactions that causes coordinated changes in gene copy number. The gene ontology (GO) and metabolic pathway analyses revealed a significant functional enrichment of genes expressed on the cell surface and involved in the interaction with the host (NC2), gibberellin biosynthesis (NC3), transmembrane nucleobase transporters (NC4) and gluconeogenesis (NC5) (**Table S7**). Altogether, GIP and giptools are validated by reproducing previously published results, and beyond that can drive new biological findings as documented by the discovery of a network of epistatic CNV interactions supporting genomic adaptation in *Candida albicans* populations under drug selection.

**Fig. 4:**
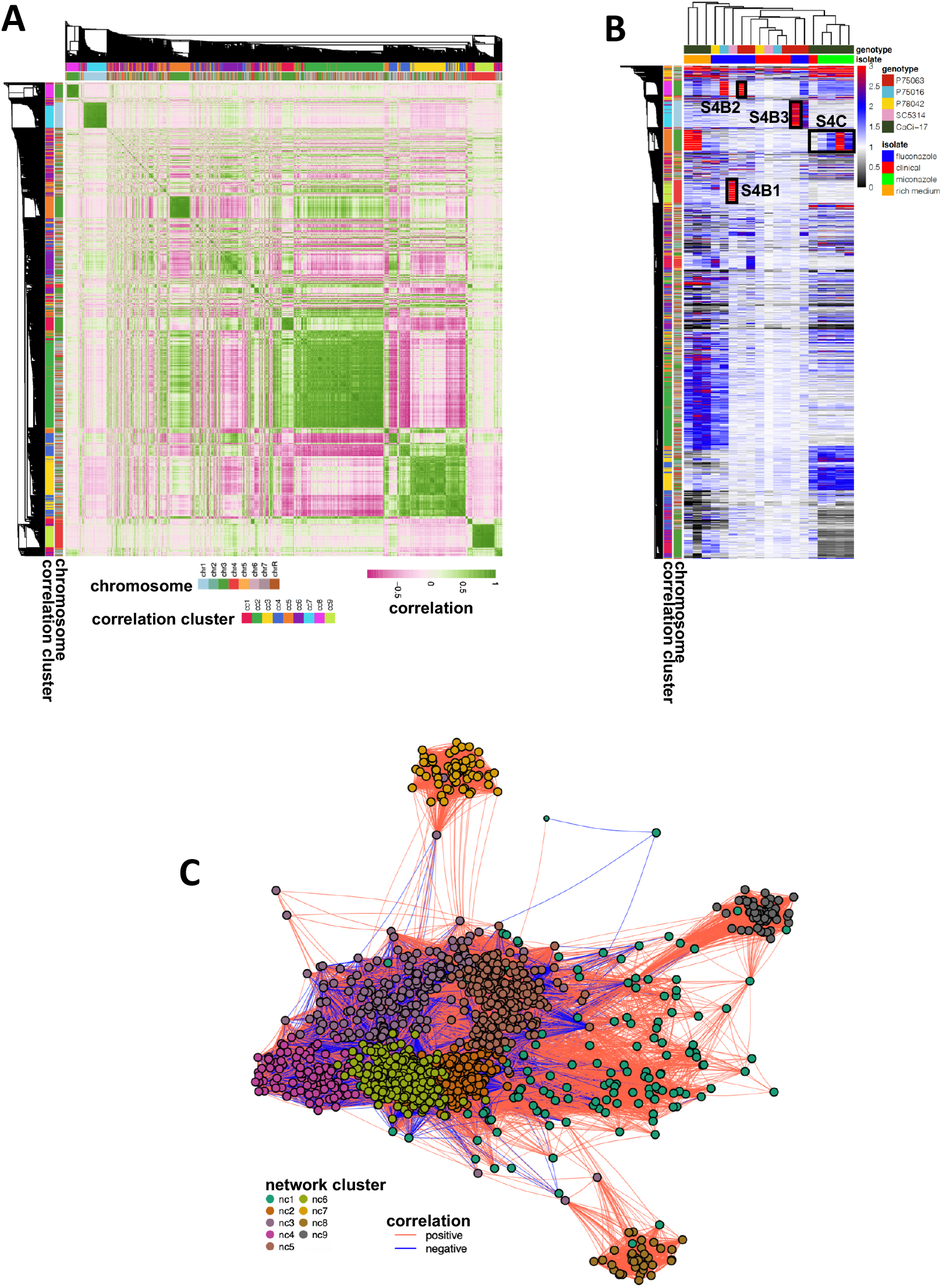
Gene CNVs interactions. (**A**) All-vs-all normalized sequencing coverage correlation heatmap. The heatmap is symmetrical along its diagonal and reports both on the rows and the columns the detected gene CNVs. The colour scale indicates with green and pink high levels of positive and negative Pearson correlation, respectively. The side ribbons demonstrate in different colours the chromosome and the correlation cluster of each gene. (**B**) Gene CNV heatmap. The columns and the rows report respectively the samples and the detected gene CNVs. The colour scale indicates the normalized sequencing coverage of the genes. To ease visualization, coverage values greater than 3 are reported as 3 (red). Black boxes highlight the genomic regions shown in **Fig. S3B** (panels 1, 2 and 3) and **Fig. S3C**. The ribbons on the left indicate the chromosome and the correlation cluster of each gene. Top ribbons indicate the genotype and the strains resulting from the different evolutionary experiments. (**C**) Gene interaction network. Nodes indicate gene CNVs. Edges reflect the absolute Pearson correlation value. The closer the nodes are, the higher is the correlation. Only significant interactions (Benjamini-Hochberg adjusted p-value < 0.01) are shown. The colour of the edges indicates in red and blue respectively positive and negative correlations. The colour of the nodes denotes the predicted network cluster for each gene.

### Exploring instability of larger genomes using cancer cell lines as a benchmark

The larger genome size, and the higher number of genes and WGS reads can represent a challenge when working with higher eukaryotes. For the purpose of comparison, the human genome is ~216 times larger than the one of *Candida albicans* we analyse in this study. Therefore, we sought to evaluate the applicability of the GIP and giptools framework to human data and utilized a panel of genomes from cancer cell lines as a test set. In our analyses we considered publicly available WGS data of the cell lines T47D, NCI_H460 and K562 (50), which respectively derive from human breast, lung and blood cancers. The karyotype analysis revealed aneuploidy for all chromosomes except chromosome 4 (**Fig. 5A**). The observed heterogeneity in read depth across chromosomes, illustrated by large interquartile range in the boxplot, suggests sub-chromosomal or episomal copy number variations, or the co-existence of karyotypically different sub-populations. Indeed, the coverage analysis confirmed the pervasive presence of CNVs both at chromosomal and sub-chromosomal levels (**Fig. S4**) with remarkable instability observed for specific chromosomes, e.g. chromosomes 6, 9, 10 and 16 (**Fig. 5B**). Overall, we detected 1,647,016 SNVs (**Supplementary Data 1**) and allele frequency shifts with respect to the reference genome, suggesting haplotype selection and the preferential expression of distinct alleles in different cell lines (**Fig. 5C** and **Fig. S5**). Furthermore, we identified repeated loss of heterozygosity events and uneven distribution of SNVs that form “patches” of high frequency correlating with chromosomal and sub-chromosomal CNVs (**Fig. 5D, Fig. 5E** and **Fig. S6**). These results identify GIP and giptools as a powerful new platform to reveal loci, genes or alleles that are under natural selection in cancer cells, thus allowing important new insight into the genetic basis of tumor development, cancer cell evolution and drug resistance.

**Fig. 5:**
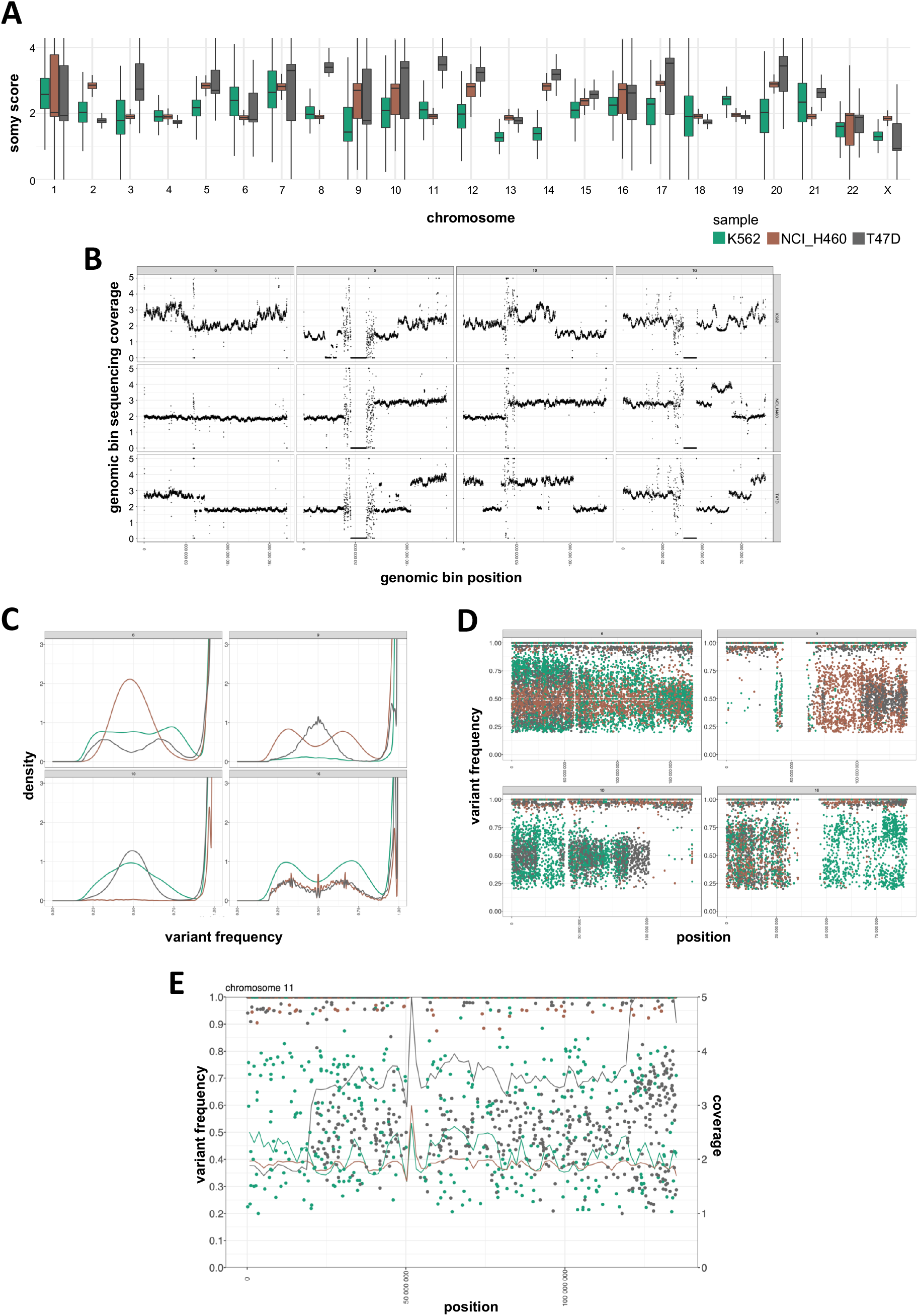
Cancer cell lines genome instability. Green, brown, and grey colours indicate respectively samples K562, NCI_H460 and T47D. (**A**) Chromosome coverage analysis. The x-axis reports the chromosomes. The y-axis reports the estimated somy score. The boxes show the somy score distributions. (**B**) Sub-chromosomal copy number variation. Dots indicate genomic bins. Different panels indicate different chromosomes. The panel columns indicate from left to right four selected chromosomes: 6, 9, 10 and 16. The panel rows show top to bottom the samples K562, NCI_H460 and T47D. The x-axis indicates the genomic position. The y-axis indicates the normalized genomic bin sequencing coverage values. Coverage values greater than 5 are reported as 5. (**C**) SNV frequency density plots. The four different panels represent different selected chromosomes: 6, 9, 10 and 16. The x-axis reports the variant allele frequency. The y-axis the estimated kernel density between 0 and 3. (**D**) SNV frequency scatter plots. The four different panels represent different selected chromosomes: 6, 9, 10 and 16. The x-axis indicates the genomic position. The y-axis indicates the variant allele frequency. (**E**) Chromosome 11 combined SNV and bin coverage plot. To ease visualization, giptools allows the simultaneous displaying of variant allele frequencies (y-axis, left) and sequencing coverage (y-axis, right). Dots represent SNVs. The lines represent the normalized bin sequencing coverage. The x-axis indicates the genomic position. Coverage values greater than 5 are shown as 5.

## Discussion

Genome instability is a key driver of evolution for microbial pathogens and cancer cells (51) and a major source of human morbidity. Here we introduce GIP and giptools, an integrated framework for the genotype profiling of biological systems exploiting genome instability for adaptation. We document the power and versatility of GIP and giptools by performing genomic screenings of three major pathogenic eukaryotes and human cancer cell lines. While originally deployed for *Leishmania* genome analysis, in this study we validate the use of our pipeline on other organisms reproducing expected results. For example, in *Candida albicans* we confirmed the CNVs correlating to drug resistance as well as a loss of heterozygosity event (**Fig. S3B-C** and **D**). Parallel to this we also show how GIP and giptools can be used for data mining and scientific discovery. New findings include (i) the discovery of the convergent amplification of erythrocyte binding proteins in *Plasmodium vivax* strains sampled from distinct geographic areas (**Fig. 3E**), (ii) the detection of a nullisomic strain (**Fig. 3F**), (iii) the identification of correlated copy number variations between genes positioned on separate chromosomes of *Candida albicans* adapting strains, and (iv) the functional association of such genes, strongly supporting a mechanism of epistatic interactions exerted through gene-dosage changes, and corroborating previous reports on adapting *Leishmania* populations (52).

Importantly, GIP and giptools overcome key limitations of current analysis tools, such as the breadth of analysis that is often limited to individual types of mutations, and the lack of genome-wide, comprehensive reports. To ease genome instability investigations our pipeline offers a single solution to karyotype, gene CNV, SNV and SV batch analyses, providing summary reports and high-quality, genome-wide visualizations. Furthermore, many current tools identify variations with respects to a reference assembly only, which leaves the between samples comparisons to external tools that need installing, may be incompatible in terms of file format, and may rely on different analytical assumptions. To address this limitation giptools enables custom sample comparisons, to explore differences and common features between genomes, and provides a vast choice of analytical tools with compatible features. Likewise, current tools are often restricted to the analysis of data from one or few species only (53–57), but are not generally applicable to different biological systems, which interferes with the investigation of genome variations across multiple species and the exploration potentially conserved genomic adaptation mechanisms. By contrast, our pipeline limits as much as possible the use of hardcoded parametrization, which could limit its use to a specific organism. Therefore, GIP’s flexible design makes it adapted for the genome analysis of both model and non-model organisms, including *Leishmania* or human.

Many current tools are further limited in software portability and reproducibility across different computer environments, which can produce faulty results calling their clinical application into question. Conversely, thanks to the Singularity implementation all required software are embedded and provided within the software container. As a consequence, users can easily recreate the same work environment just by downloading the pipeline container, and reproduce exactly the same publication-quality plots and tables presented in this study. Lastly, one more common limitation is posed by software scalability. In the WGS domain, with the rapid increase new samples made available and the enormous amount of data generated in each sequencing run, the CPU and memory resources of local workstation risk to quickly become inadequate for data analysis. Therefore, it is paramount that WGS tools are implemented to run on high-performance computing (HPC) clusters and feature remote cloud computing solutions. Because of its Nextflow implementation GIP can be executed on a local machine, on cluster resource manager or the cloud. GIP can be applied on individual samples and without additional effort on large WGS data sets for batch computation as shown for the 222 *Plasmodium vivax* genomes.

These results well illustrate how GIP and giptools can be applied to perform extended genomic analyses in different biological systems and drive biomedical discovery. To conclude, we believe that GIP and giptools represent a step forward toward reproducible research in genomics, and provide a robust computational framework to study how microbes and tumor cells harness genome instability for environmental adaptation and fitness gain.

## Methods

### GIP and giptools

All results presented in this study were generated using GIP and giptools version 1.0.9. GIP code is maintained and freely distributed at the github page: https://github.com/giovannibussotti/GIP. giptools container is accessible from the Singularity cloud at https://cloud.sylabs.io/library/giovannibussotti/default/giptools. The GIP configuration files (**Supplementary Data 2**) and the giptools command options used to generate all results (**Supplementary Data 3**) are provided. The full documentation of GIP and giptools including a description of all options is available from https://gip.readthedocs.io/en/latest/.

### Read alignment

WGS reads were downloaded from the Sequence Read Archive (SRA) (58) and the European Nucleotide Archive (ENA) (59) repositories and the Encyclopedia of DNA Elements (ENCODE) dashboard (60) (**Table S3**). For *Leishmania infantum* the GCA_900500625 genome reference and gene annotations available from the ENSEMBL protists (61) server (release-48) were used. For *Candida albicans* the assembly 21 of the SC5314 strain genome reference and gene annotations available from the *Candida* Genome Database (CGD) (62) were used. For *Plasmodium vivax* the P01 reference genome and gene annotations available from PlasmoDB (63) (release-50) were used. For the cancer cell lines the human genome GRCh38 primary assembly and gene annotations available from ENSEMBL (release-102) were used. The repetitive elements of reference genomes were soft-masked by GIP using Red (64). WGS reads were mapped by GIP using BWA-mem (version 0.7.17) (65,66) run with option -M to label shorter split hits as secondary. Then the alignment files were sorted, indexed and reformatted by GIP using Samtools (version 1.8) (67). Finally, read duplicates were removed by GIP using Picard MarkDuplicates (http://broadinstitute.github.io/picard) (version 2.18.9) with the option “VALIDATION_STRINGENCY=LENIENT.” In the four considered datasets, WGS reads were aligned against full assemblies, including unsorted contigs if present. However just the canonical assembled chromosomes were considered for all downstream analyses (‘chrs’ option, **Supplementary Data 2**). A minimum read alignment MAPQ score was adopted to select genes for cluster analysis, and to call for SNVs and SVs (‘MAPQ’ option, **Supplementary Data 2**). Altogether a total of 6,306,951,266 reads were aligned to the respective reference genomes. The ‘giptools overview’ module was run to gather the alignment statistics as estimated by Picard CollectAlignmentSummaryMetrics (**Table S8**).

### Genomic bins and genes quantification

GIP was used to evaluate the mean sequencing coverage and the mean read MAPQ of genomic bins and genes. For genomic bins, GIP partitioned the input genomes into adjacent windows of user defined lengths (‘binSize’ option, **Supplementary Data 2**). The coverage GC-content score bias was corrected (‘CGcorrect’ option, **Supplementary Data 2**) fitting a LOESS regression with a 5-fold cross validation to optimize the model span parameter. A larger window length was utilized to bin the reference genomes for Circos plot representations (‘binSizeCircos’ option, **Supplementary Data 2**). In **Fig. 1** (“Genomic bins” and “Gene CNVs” plots), **Fig. 2, Fig. S1, Fig. S2** and **Fig. 4** bins and genes coverage scores were normalized by median chromosome coverage to highlight amplifications or depletions with respect to the chromosome copy number. In **Fig. 1** (“Structural variants” plot), **Fig. S3B-C, Fig. 3E-G**, and **Fig. S4A** bins and genes coverage scores were normalized by median genome coverage to account for sequencing library size differences. GIP evaluated statistically significant copy number variant bins and genes (**Fig. 1** “Genomic bins” and “Gene CNVs” plots) using a p-value threshold of 0.001 (‘covPerBinSigOPT’ and ‘covPerGeSigOPT’ options, **Supplementary Data 2**). Estimated p-values for bins and genes CNVs were corrected for multiple testing using the Benjamini – Yekutieli (‘--padjust BY’) and the Benjamini – Hochberg (‘--padjust BH’) methods. The somy scores shown in **Fig. 1** (karyotype plot) and **Fig. S4B** were computed multiplying the median genome coverage normalized bin coverage by 2. GIP enabled the CNV analysis of genes sharing high sequence identity by clustering the nucleotide sequences of the genes with low mean MAPQ score into groups with cd-hit-est (version 4.8.1) (68) with options ‘-s 0.9 -c 0.9 - r 0 -d 0 -g 1’. Then for each gene cluster GIP computed the mean gene coverage normalized by median chromosome coverage (**Fig. 2E, Fig S2B**).

### Gene ontology and metabolic pathway enrichment

The FungiDB online tool (Release 52, 20 May 2021) (69) was used to evaluate the functional enrichment of network clusters genes. For the gene ontology analysis, the biological process (BP), molecular function (MF) and cellular compartment (CC) terms enrichments were tested, considering both computed and curated evidences and a p-value cutoff of 0.05. For the metabolic pathway enrichment, both KEGG (70) and MetaCyc (71) pathway sources were considered with a p-value cutoff of 0.05. Terms and pathways with Benjamini – Hochberg adjusted p-values < 0.05 were considered statistically significant.

### Sequencing coverage density estimates

GIP was used to convert the read alignment files (.bam format) in binary data files reflecting sequencing coverage (.bigWig format). The coverage file were produced using bamCoverage from the deepTools2 suite (72) (version 3.5.1) with options “--normalizeUsing RPKM --ignoreDuplicates --binSize 10 --smoothLength 30” (‘bigWigOPT’ option, **Supplementary Data 2**). The coverage track of sample PD0689_C was visualized with IGV using the ‘Bar Chart’, ‘Autoscale’ and windowing function ‘Mean’ options.

### Single-nucleotide variant analysis

GIP was used to call SNVs using Freebayes (version 1.3.2) (‘freebayesOPT’ option, **Supplementary Data 2**) and filter its output (‘filterFreebayesOPT’ option, **Supplementary Data 2**). Filters included the minimum allele frequency (‘--minFreq’), the minimum number of reads supporting the alternative alleles (‘-- minAO’) and minimum mean mapping quality of for the reads supporting the reference (‘-- minMQMR’) or the alternative allele (‘--minMQM’). A higher number of reads supporting the variants was requested for predictions positioned inside simple repeats of the same nucleotide (homopolymers) (‘--minAOhomopolymer’). The homopolymers were defined as the DNA region spanning ±5 bases from the SNV (‘--contextSpan 5’), with over 40% of identical nucleotides (--homopolymerFreq 0.4’). Further, GIP discarded SNVs with sequencing coverage above or below 4 median absolute deviations (MADs) from the median chromosome coverage (‘--MADrange’). snpEff (version 4.3t) (73) was used to predict the impact of SNVs on coding sequence. The predicted effects that GIP considered synonymous mutations are: “synonymous_variant”, “stop_retained_variant” and “start_retained”. The predicted effects that GIP considered non-synonymous mutations are: “missense_variant”, “start_lost”, “stop_gained”, “stop_lost” and “coding_sequence_variant”. The phylogenetic tree was computed by the giptools module ‘phylogeny’ using IQtree2 (version 2.1.2) (74,75) with options ‘--seqtype DNA --alrt 1000 -B 1000’. The Venn-diagram comparison considered the strains QS0044_C, QS0001_C, QS0037_C, QS0016_C and SGH_358 that were sampled from different locations in Ethiopia, respectively Habala, Badowacho, Arbaminch, Hawassa and Jimma. The strains were selected to have comparable average genome coverage (23). To infer the tree GIP considered the set of filtered SNV and adopted the IUPAC ambiguous notation for the positions with allele frequency below 70%. The tree was visualized by giptools using the ggtree R-package (76).

### Analysis of structural variants

GIP was used to detect structural variants including insertions, tandem duplications, deletions, inversions and break-end translocations with DELLY (version 0.8.7) (11). To reduce incorrect predictions the DELLY output was additionally filtered (‘filterDellyOPT’ option, **Supplementary Data 2**). GIP discarded poor predictions with DELLY label “LowQual” (‘--rmLowQual’) and low median MAPQ score of mapping reads (‘--minMAPQ’). SVs positioned in proximity of chromosome ends were removed (-- chrEndFilter) to limit false predictions caused by potential misassembled regions close to the telomeric ends. To ease visualization and limit the analysis only to best supported SVs GIP limited the output only to the top predictions (‘--topHqPercentIns’, ‘--topHqPercentDel’, ‘-- topHqPercentDup’ and ‘--topHqPercentInv’) based on the SV support score as in **Formula 1**, where *DV*, *DR*, *RV*, and *RR* are respectively the number of high-quality variant pairs, reference pairs, variant junction reads, and reference junction reads.

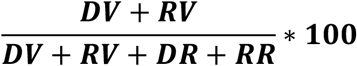

**Formula 1:** SV support score.

The predicted structural variants were represented with Circos (version 0.69-9) (77).

## Data and materials availability

All data is available in the main text or the supplementary materials.

## Author contributions

Conceptualization, G.B. and G.F.S.; Methodology, G.B. and G.F.S.; Software, G.B.; Formal Analysis, G.B., and G.F.S.; Investigation, G.B. and G.F.S.; Writing – Original Draft, G.B. and G.F.S.; Writing – Review & Editing, G.B. and G.F.S.; Supervision, G.B. and G.F.S.; Funding Acquisition, G.F.S. and G.B.

## Supplementary Tables

**Table S1:** Genomic bin coverage ratio of sample LIPA83 over ZK43.

**Table S2:** Gene coverage ratio of sample LIPA83 over ZK43.

**Table S3:** Sample information.

**Table S4:** *Candida albicans* gene CNVs.

**Table S5:** Gene correlation clusters in *Candida albicans*.

**Table S6:** Network analysis of gene CNVs in *Candida albicans*.

**Table S7:** Gene ontology and metabolic pathway enrichment analyses.

Footnote: Terms and pathways with Benjamini – Hochberg adjusted p-values < 0.05 are labeled in red.

**Table S8:** Mapping statistics.

## Supplementary Data

**Supplementary Data 1:** Cancer cell lines SNVs.

**Supplementary Data 2:** GIP configuration files.

**Supplementary Data 3:** giptools commands.

## Supplementary figures

**Fig. S1:**
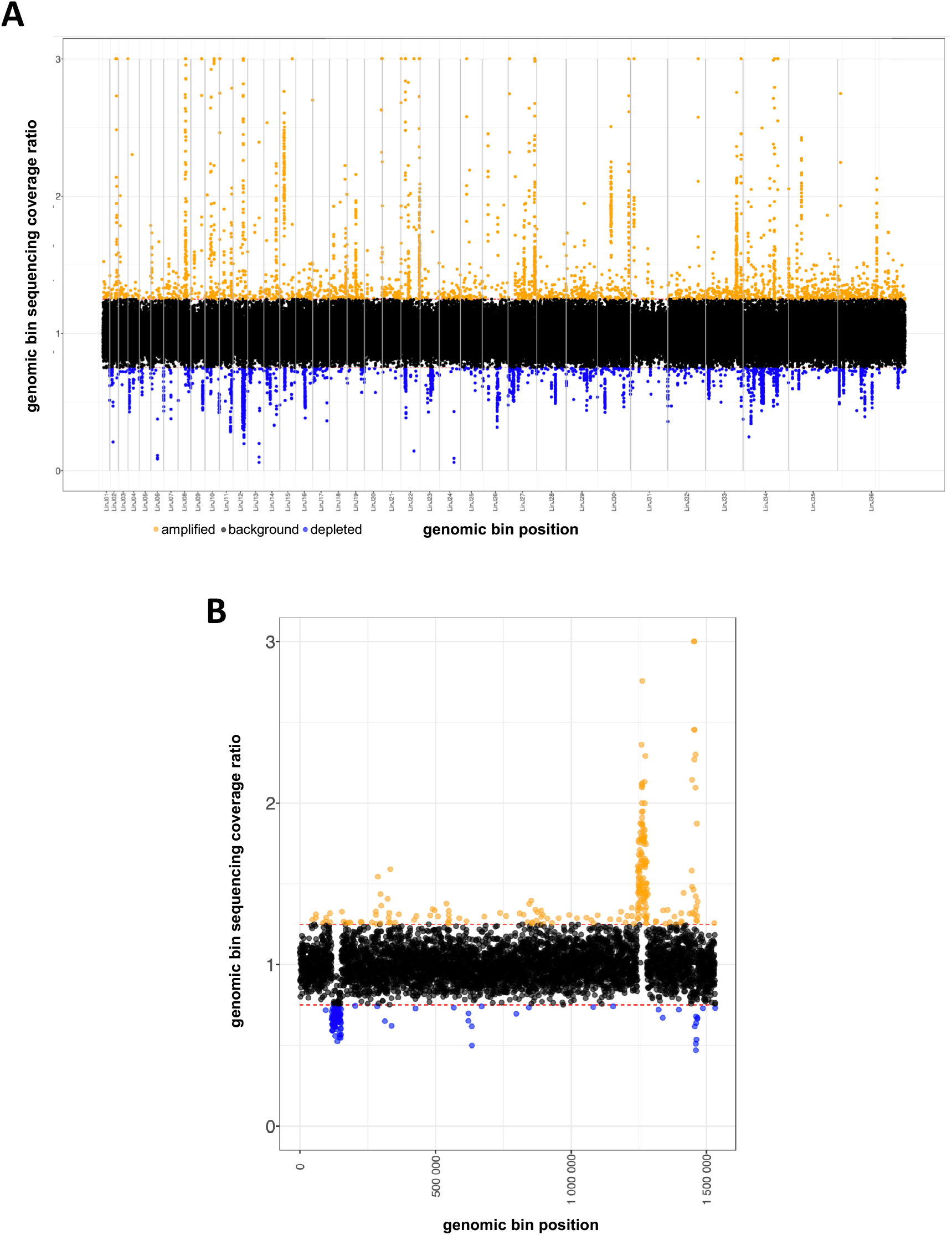
Bin coverage ratio visualizations. The figure shows alternative representations produced for the genomic bin normalized sequencing coverage ratio of samples ZK43 and LIPA83. Same layout as **Fig. 2B**. (**A**) Whole genome overview. (**B**) All the individual chromosomes separately. The plot shows the example of chromosome 33.

**Fig. S2:**
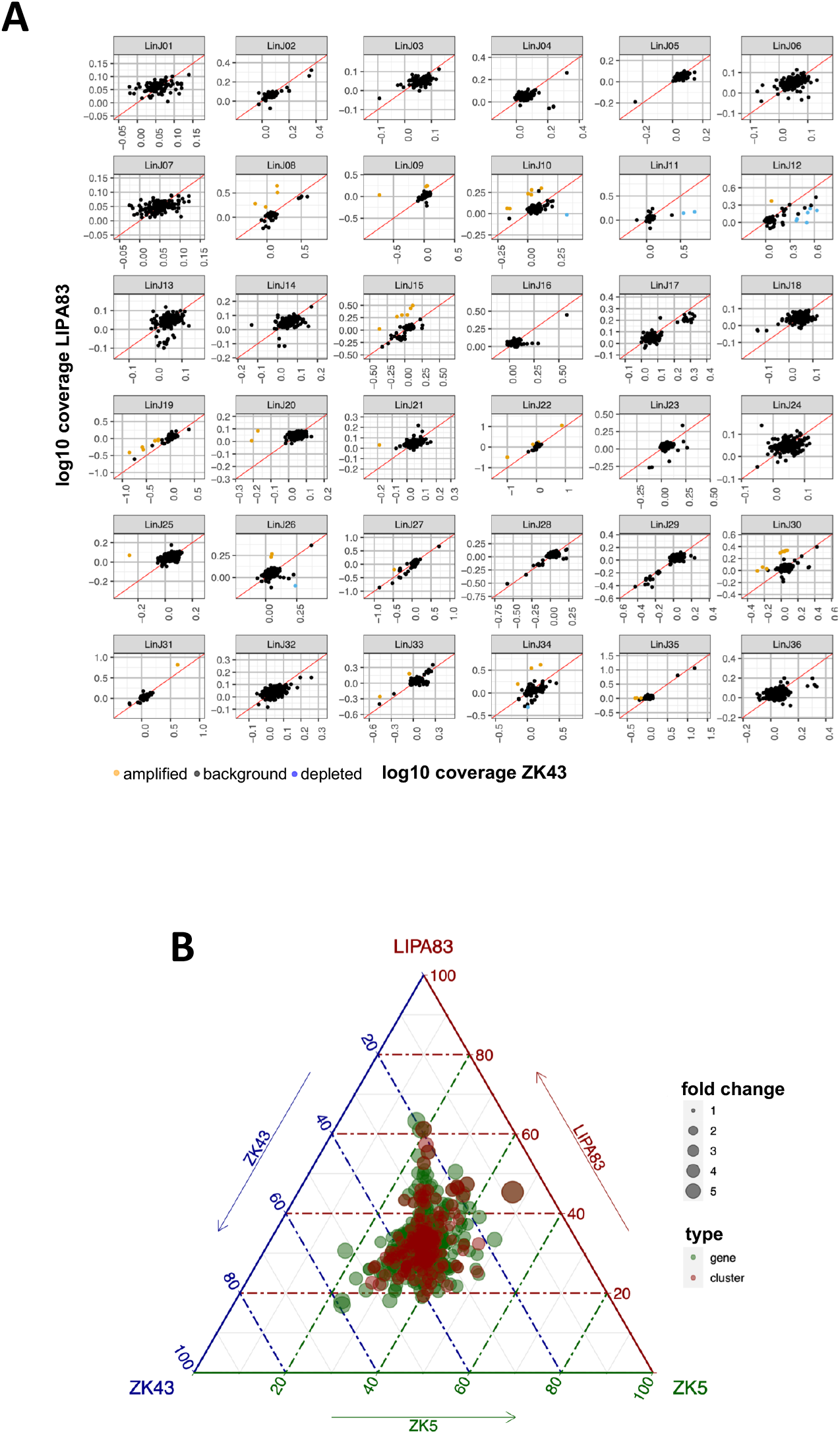
Two-ways and three-ways gene coverage comparisons. (**A**) Log10 normalized sequencing coverage comparison of genes in ZK43 (x-axis) and LIPA83 (y-axis). Same layout a **Fig. 2D** but representing the different chromosomes in individual panels. (**B**) Ternary plot showing the relative abundance of genes (green dots) and gene clusters (red dots) in samples LIPA83, ZK43 and ZK28. The axes report the fraction of the genes and genes clusters normalized sequencing coverage in the three strains. Each given point in the plot adding up to 100. Genes with equal copy number are shown in the center of the ternary plot, while copy number variations are visualized by shifts of the dots out of the center. The dots size reflects the fold change variation between the maximum and the minimum observed normalized coverage value.

**Fig. S3:**
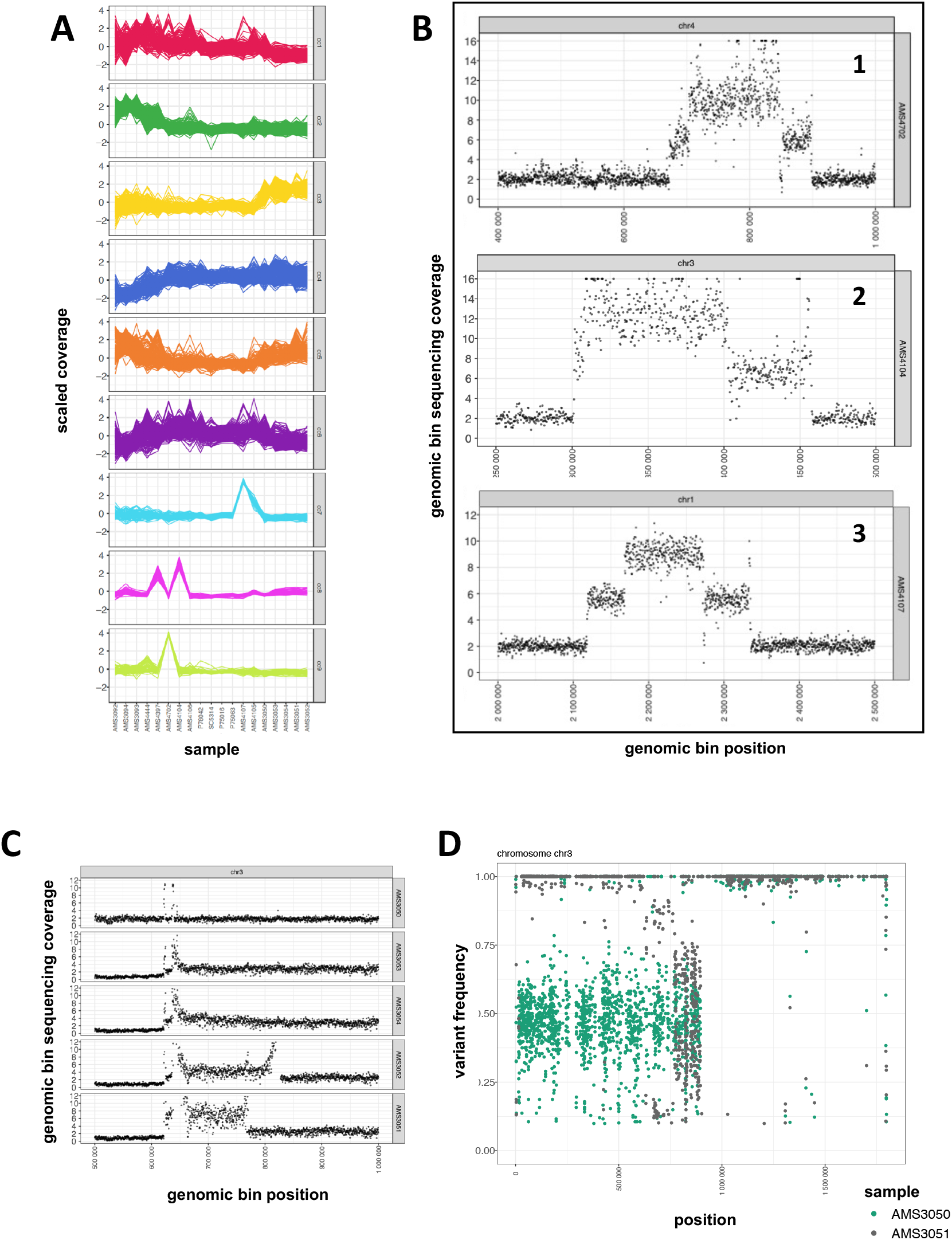
Copy number variations of evolutionary adapted *Candida albicans* strains. (**A**) Gene CNV correlation clusters. The x and y axes indicate respectively the samples and the scaled normalized sequencing coverage of the genes in each cluster. Different panels indicate different correlation clusters. (**B**) Scatterplots showing the bin normalized sequencing coverage (y-axis) of three genomic regions (chr4:400,000-1,000,000; chr3:1,250,000-1,500,000; chr1:2,000,000-2,500,000) in three different samples AMS4702, AMS4104 and AMS4107. The x-axis shows the genomic position. Dots represent genomic bins. (**C**) Scatterplot showing the bin normalized sequencing coverage of a genomic region (chr3:500,000-1,000,000) in five different samples represented as separate panels. From top to bottom the samples are: AMS3050, AMS3053, AMS3054, AMS3052 and AMS3051. The x and y axes show respectively the genomic position and the normalized sequencing coverage. Dots represent bins. (**D**) Comparative analysis of chromosome 3 SNVs. AMS3050 SNVs are shown in green, while AMS3051 SNVs are shown in grey. The x and the y axes show respectively the genomic position and the variant allele frequency.

**Fig. S4:**
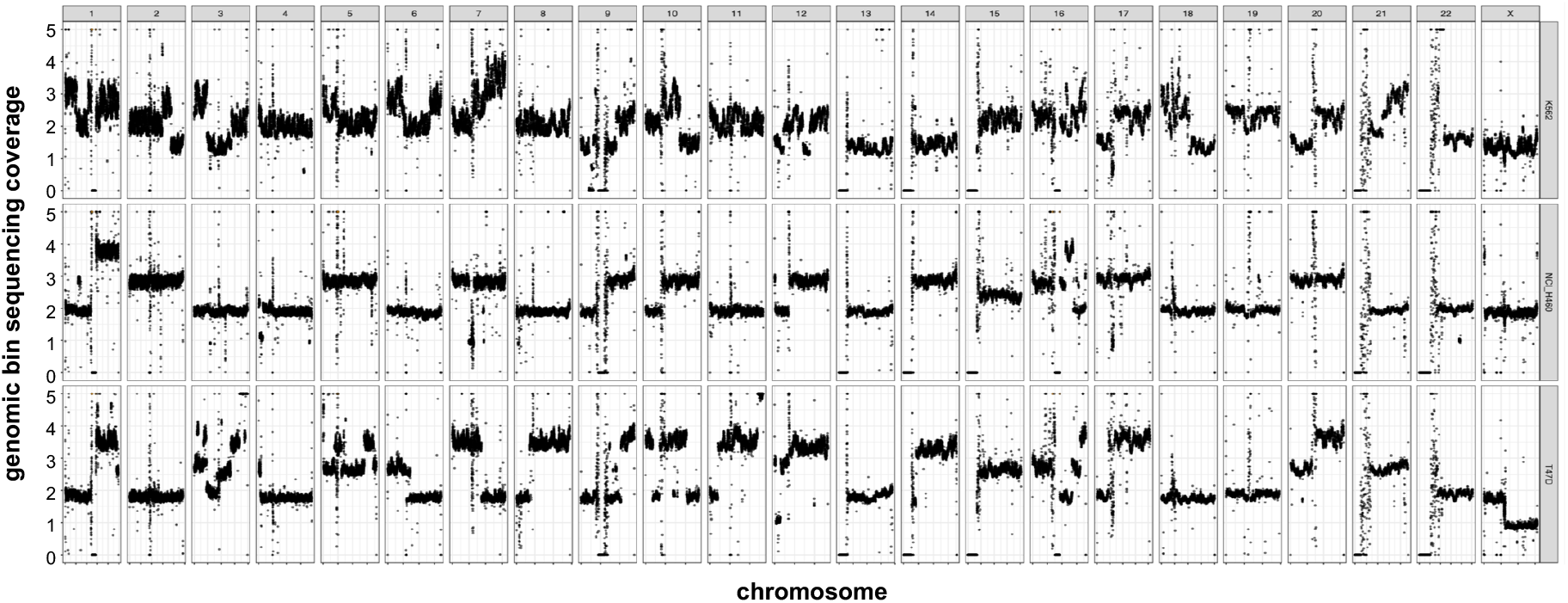
Genomic coverage analysis of three cancer cell lines. The x-axis indicates the genomic position. The y-axis indicates the normalized genomic bin sequencing coverage values. Dots demonstrate the genomic bins. To ease visualization, coverage values greater than 5 are reported as 5. Different panels show different chromosomes. The three panel rows indicate top to bottom the following samples: K562, NCI_H460 and T47D.

**Fig. S5:**
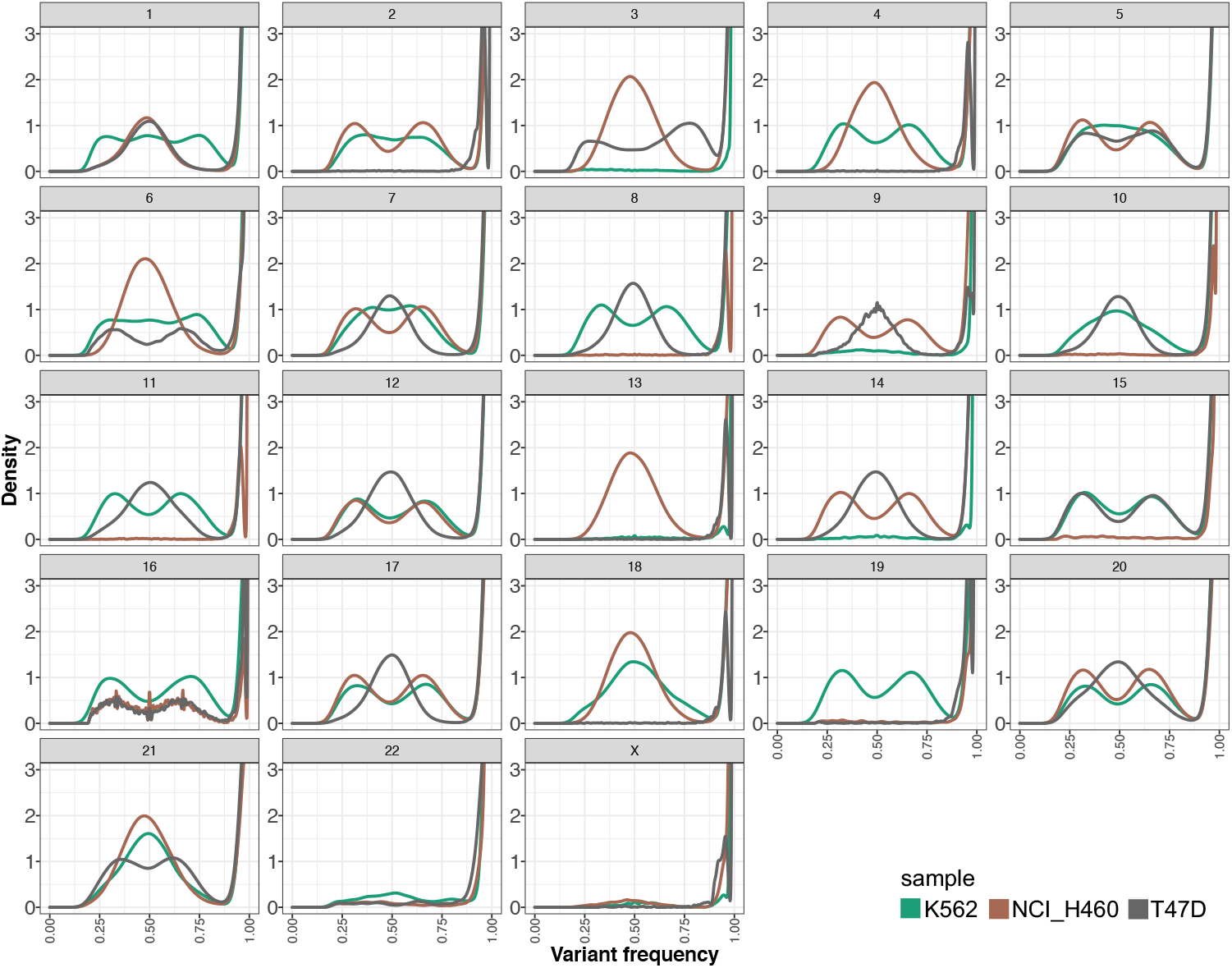
SNV frequency density plot. The x-axis shows the variant allele frequency. The y-axis the estimated kernel density between 0 and 3. Green, brown, and grey slopes indicate respectively samples K562, NCI_H460 and T47D.

**Fig. S6:**
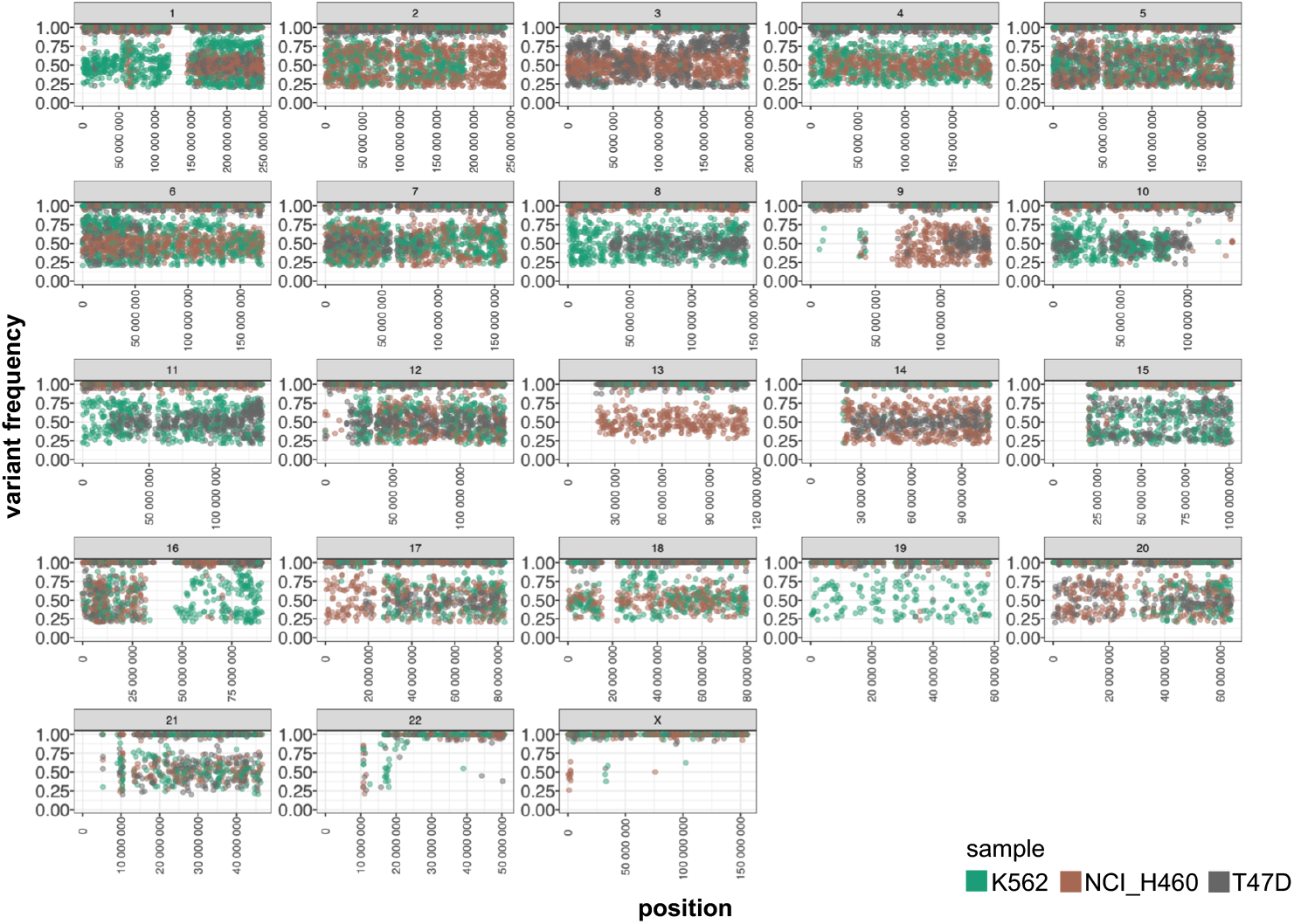
SNV heterogeneity of cancer cell lines. The x-axis indicates the genomic position. The y-axis indicates the variant allele frequency. The dots indicate SNVs and they are coloured according to the sample. K562, green; NCI_H460, brown; T47D, grey. Different panels show different chromosomes.

