## Supplementary Data 2 for "GIP: An open-source computational pipeline for mapping genomic instability from protists to cancer cells"

**Supplementary Data 2. GIP configuration file: *Leishmania* *infantum***

params{

MAPQ	= 50

delDup = true

CGcorrect = true

chrs = "all"

chrPlotYlim = "0 8"

binPlotYlim = "0 3"

binOverviewSize = "400 1000"

customCoverageLimits = "1.5 0.5"

binSize = 300

covPerBinSigOPT = "--minLen 0 --pThresh 0.001 --padjust BY"

covPerGeSigOPT = "--pThresh 0.001 --padjust BH --minLen 0"

covPerGeRepeatRange = 1000

freebayesOPT = "--read-indel-limit 1 --read-mismatch-limit 3 --read-snp-limit 3 --min-alternate-fraction 0.05 --min-base-quality 5 --min-alternate-count 2 --pooled-continuous"

filterFreebayesOPT = "--minFreq 0.1 --maxFreq 1 --minAO 2 --minAOhomopolymer 20 --contextSpan 5 --homopolymerFreq 0.4 --minMQMR 20 --minMQM 20 --MADrange 4"

filterDellyOPT="--rmLowQual --chrEndFilter 100 --minMAPQ 50 --topHqPercentBnd 150 --topHqPercentIns 150 --topHqPercentDel 150 --topHqPercentDup 150 --topHqPercentInv 150"

binSizeCircos = 25000

bigWigOPT = "--binSize 10 --smoothLength 30"

}

process{

executor='slurm'

clusterOptions='--qos normal'

container='/pasteur/zeus/projets/p01/BioIT/Giovanni/apps/installation/GIP/version_1.0.9/giptools'

errorStrategy='terminate'

scratch=false

cache=true

echo=false

stageInMode='symlink'

time='24h'

cpus=1

memory='30000'

withName: prepareGenome {

cpus=2

}

withName: map {

cpus=4

}

withName: covPerChr {

memory='30000'

}

withName: covPerBin {

memory='30000'

}

withName: mappingStats{

memory='10000'

}

withName: covPerGe {

memory='30000'

}

withName: freebayes {

memory='40000'

}

withName: snpEff {

memory='30000'

}

withName: delly {

memory='30000'

}

withName: bigWigGenomeCov {

memory='10000'

}

withName: report {

memory='10000'

}

}

singularity{

enabled	= true

autoMounts = true

runOptions = '--bind /pasteur'

}

**Supplementary Data 2. GIP configuration file: *Plasmodium vivax***

params{

MAPQ	= 5

delDup = true

CGcorrect = true

chrs = "\"chr01 chr02 chr03 chr04 chr05 chr06 chr07 chr08 chr09 chr10 chr11 chr12 chr13 chr14\""

chrPlotYlim = "0 8"

binPlotYlim = "0 3"

binOverviewSize = "400 1000"

customCoverageLimits = "1.5 0.5"

binSize = 300

covPerBinSigOPT = "--minLen 0 --pThresh 0.001 --padjust BY"

covPerGeSigOPT = "--pThresh 0.001 --padjust BH --minLen 0"

covPerGeRepeatRange = 1000

freebayesOPT = "--read-indel-limit 0 --read-mismatch-limit 3 --read-snp-limit 3 --min-alternate-fraction 0.1 --min-base-quality 27 --min-alternate-count 2 --min-coverage 5 --pooled-continuous --min-mapping-quality 27"

filterFreebayesOPT = "--minFreq 0.1 --maxFreq 1 --minAO 2 --minAOhomopolymer 20 --contextSpan 5 --homopolymerFreq 0.4 --minMQMR 20 --minMQM 20 --MADrange 4"

filterDellyOPT="--rmLowQual --chrEndFilter 1000 --minMAPQ 10 --topHqPercentBnd 150 --topHqPercentIns 150 --topHqPercentDel 150 --topHqPercentDup 150 --topHqPercentInv 150"

binSizeCircos = 25000

bigWigOPT = "--normalizeUsing RPKM --ignoreDuplicates --binSize 10 --smoothLength 30"

}

process{

executor='slurm'

clusterOptions='--qos normal'

container='/pasteur/zeus/projets/p01/BioIT/Giovanni/apps/installation/GIP/version_1.0.9/giptools'

errorStrategy='terminate'

scratch=false

cache=true

echo=false

stageInMode='symlink'

time='24h'

cpus=1

memory='30000'

withName: prepareGenome {

cpus=2

}

withName: map {

cpus=4

}

withName: covPerChr {

memory='30000'

}

withName: covPerBin {

memory='30000'

}

withName: mappingStats{

memory='10000'

}

withName: covPerGe {

memory='30000'

}

withName: freebayes {

memory='40000'

}

withName: snpEff {

memory='30000'

}

withName: delly {

memory='30000'

}

withName: bigWigGenomeCov {

memory='10000'

}

withName: report {

memory='10000'

}

}

singularity{

enabled	= true

autoMounts = true

runOptions = '--bind /pasteur'

}

**Supplementary Data 2. GIP configuration file: *Candida albicans***

params{

MAPQ	= 5

delDup = true

CGcorrect = true

chrs = "all"

chrPlotYlim = "0 8"

binPlotYlim = "0 3"

binOverviewSize = "400 1000"

customCoverageLimits = "1.5 0.5"

binSize = 300

covPerBinSigOPT = "--minLen 0 --pThresh 0.001 --padjust BY"

covPerGeSigOPT = "--pThresh 0.001 --padjust BH --minLen 0"

covPerGeRepeatRange = 1000

freebayesOPT = "--read-indel-limit 1 --read-mismatch-limit 3 --read-snp-limit 3 --min-alternate-fraction 0.05 --min-base-quality 5 --min-alternate-count 2 --pooled-continuous"

filterFreebayesOPT = "--minFreq 0.1 --maxFreq 1 --minAO 2 --minAOhomopolymer 20 --contextSpan 5 --homopolymerFreq 0.4 --minMQMR 20 --minMQM 20 --MADrange 4 --randomSNVtoShow 50000"

filterDellyOPT="--rmLowQual --chrEndFilter 100 --minMAPQ 50 --topHqPercentBnd 150 --topHqPercentIns 150 --topHqPercentDel 150 --topHqPercentDup 150 --topHqPercentInv 150"

binSizeCircos = 25000

bigWigOPT = "--binSize 10 --smoothLength 30"

}

process{

executor='slurm'

clusterOptions='--qos normal'

container='/pasteur/zeus/projets/p01/BioIT/Giovanni/apps/installation/GIP/version_1.0.9/giptools'

errorStrategy='terminate'

scratch=false

cache=true

echo=false

stageInMode='symlink'

time='24h'

cpus=1

memory='30000'

withName: prepareGenome {

cpus=2

}

withName: map {

cpus=4

}

withName: covPerChr {

memory='30000'

}

withName: covPerBin {

memory='30000'

}

withName: mappingStats{

memory='10000'

}

withName: covPerGe {

memory='30000'

}

withName: freebayes {

memory='40000'

}

withName: snpEff {

memory='30000'

}

withName: delly {

memory='30000'

}

withName: bigWigGenomeCov {

memory='10000'

}

withName: report {

memory='10000'

}

}

singularity{

enabled	= true

autoMounts = true

runOptions = '--bind /pasteur'

}

**Supplementary Data 2. GIP configuration file: *Homo sapiens***

params{

MAPQ	= 5

delDup = true

CGcorrect = false

chrs = "\"1 2 3 4 5 6 7 8 9 10 11 12 13 14 15 16 17 18 19 20 21 22 X\""

chrPlotYlim = "0 8"

binPlotYlim = "0 3"

binOverviewSize = "400 1000"

customCoverageLimits = "1.5 0.5"

binSize = 50000

covPerBinSigOPT = "--minLen 0 --pThresh 0.001 --padjust BY"

covPerGeSigOPT = "--pThresh 0.001 --padjust BH --minLen 0"

covPerGeRepeatRange = 1000

freebayesOPT = "--read-indel-limit 0 --read-mismatch-limit 2 --read-snp-limit 2 --min-alternate-fraction 0.2 --min-base-quality 20 --min-coverage 10 --min-alternate-count 5 --pooled-continuous --min-mapping-quality 27"

filterFreebayesOPT = "--minFreq 0.20 --maxFreq 1 --minAO 5 --minAOhomopolymer 30 --contextSpan 5 --homopolymerFreq 0.4 --minMQMR 20 --minMQM 20 --MADrange 4 --randomSNVtoShow 50000"

filterDellyOPT="--rmLowQual --chrEndFilter 10000 --minMAPQ 50 --topHqPercentBnd 150 --topHqPercentIns 150 --topHqPercentDel 150 --topHqPercentDup 150 --topHqPercentInv 150"

binSizeCircos = 2500000

bigWigOPT = "--binSize 10 --smoothLength 30"

}

process{

executor='slurm'

clusterOptions='-p hubbioit --qos hubbioit'

container='/pasteur/zeus/projets/p01/BioIT/Giovanni/apps/installation/GIP/version_1.0.9/giptools'

errorStrategy='terminate'

scratch=false

cache=true

echo=false

stageInMode='symlink'

//time='20h'

cpus=1

memory='30 GB'

withName: prepareGenome {

cpus=6

}

withName: map {

memory='50 GB'

cpus=10

}

withName: covPerChr {

memory='40 GB'

}

withName: covPerBin {

memory='30 GB'

}

withName: mappingStats{

memory='10 GB'

}

withName: covPerGe {

memory='30 GB'

}

withName: freebayes {

memory='50 GB'

}

withName: snpEff {

memory='30 GB'

}

withName: delly {

memory='30 GB'

}

withName: bigWigGenomeCov {

memory='10 GB'

}

withName: report {

memory='10 GB'

}

}

singularity{

enabled	= true

autoMounts = true

runOptions = '--bind /pasteur'

}
