## Supplementary Data 3 for "GIP: An open-source computational pipeline for mapping genomic instability from protists to cancer cells"

**Supplementary Data 3. giptools commands**

**Setup**

#!/bin/bash

module load singularity/3.6.4

mkdir -p tmpDir

export TMPDIR=$PWD/tmpDir

container=/pasteur/zeus/projets/p01/BioIT/Giovanni/apps/installation/GIP/version_1.0.9/giptools

giptools="singularity run -B /pasteur $container"

***Leishmania* *infantum* commands**

#Fig.2A

$giptools binDensity

#Fig.2B - Fig.S1 - Table S1

$giptools binCNV --samples ZK43 LIPA83 --highLowRatio 1.25 0.75

#Fig.2C

$giptools ternaryBin --samples ZK43 LIPA83 ZK28

#Fig.2D - Fig.2E - Fig.S2A - Table S2

$giptools geCNV --samples ZK43 LIPA83

#Fig.S2B

$giptools ternary --samples ZK43 LIPA83 ZK5

#Table S8

$giptools overview --noBin

***Plasmodium vivax* commands**

#Fig.3A - Fig.3B

$giptools phylogeny --iqtreeOpts \"--seqtype DNA --alrt 1000 -B 1000 -T 10\"

$giptools geInteraction --cnvPlotDim 11 11 --samplesList sampleList --clusteringMethod ward.D2 --rmNotSigGenes --kmeansClusters 2 --heatmapType flatten

$giptools convergentCNV --treeTip country --hideXlabs --plotDim 5 5 --branchLen branch.length --hideHeatmap

$giptools genomeDistance --PCAlabel none --phylogenyDistance gipOut/sampleComparison/phylogeny.treefile.distMat.gz

#Fig.3C - Fig.3D

$giptools SNV --samples QS0044_C QS0001_C QS0037_C QS0016_C SGH_358

#Fig.3F is generated visualizing with IGV the coverage density track denerated by GIP

#Fig.3E - Fig.3G

$giptools panel --samplesList sampleList --panel panel.tsv --contrast Ethiopia Cambodia --varPlotDim 5 10 --covPlotDim 5 10

#Table S8

$giptools overview --noBin

***Candida albicans* commands**

#Fig.4 - Fig.S3A - Table S4 - Table S5 - Table S6

$giptools geInteraction --chrs chr1 chr2 chr3 chr4 chr5 chr6 chr7 chrR --samplesList sampleList --clusteringMethod single --heatmapType saturated --minMaxCov 2 0.5 --clMaxSDdist 1.5

#Fig.S3A

$giptools geInteraction --chrs chr1 chr2 chr3 chr4 chr5 chr6 chr7 chrR --samplesList sampleList --clusteringMethod single --heatmapType saturated --minMaxCov 2 0.5 --clMaxSDdist 1.5 --corPlotDim 15 5

#Fig.S3B(1)

$giptools overview --chrs chr1 chr3 chr4 --samples AMS4702 SC5314 --ylim 16 --ylimInt 2 --highLowCovThresh 10000000 -1 --minDelta 1 --regions regions2

#Fig.S3B(2)

$giptools overview --chrs chr1 chr3 chr4 --samples AMS4397 AMS4104 --ylim 16 --ylimInt 2 --highLowCovThresh 10000000 -1 --minDelta 1 --regions regions2

#Fig.S3B(3)

$giptools overview --chrs chr1 chr3 chr4 --samples AMS4107 P75063 --ylim 12 --ylimInt 2 --highLowCovThresh 10000000 -1 --minDelta 1 --regions regions2

#Fig.S3C

$giptools overview --chrs chr3 --samples AMS3050 AMS3053 AMS3054 AMS3052 AMS3051 --ylim 12 --ylimInt 2 --highLowCovThresh 10000000 -1 --minDelta 1 --regions regions

#Fig.S3D

$giptools SNV --chrs chr1 chr3 --samples AMS3050 AMS3051

#Table S8

$giptools overview --noBin

***Homo sapiens* commands**

#Fig.5A

$giptools karyotype --ylim 0 7

#Fig.5B

$giptools overview --highLowCovThresh 10000 0 --chrs 6 9 10 16

#Fig.5C-D

$giptools SNV --randomSNVtoShow 50000 --showCoverage --coordCartesianYlim 3 --chrs 6 9 10 16

#Fig.S4 - Table S8

$giptools overview --highLowCovThresh 10000 0

#Fig.S5 - Fig.S6 - Fig. 5E

$giptools SNV --randomSNVtoShow 50000 --showCoverage --coordCartesianYlim 3
